# Label-free digital holo-tomographic microscopy reveals virus-induced cytopathic effects in live cells

**DOI:** 10.1101/401075

**Authors:** Artur Yakimovich, Robert Witte, Vardan Andriasyan, Fanny Georgi, Urs F. Grebera

**Affiliations:** Department of Molecular Life Sciences, University of Zurich, Winterthurerstrasse 190, 8057 Zurich, Switzerland; MRC Laboratory for Molecular Cell Biology, University College London, Gower St, London WC1E 6BT, UK

**Keywords:** Live cell Imaging, Virus Infection, Holography, Label-free microscopy

## Abstract

Cytopathic effects (CPEs) are a hallmark of infections. CPEs can be observed by phase contrast or fluorescence light microscopy, albeit at the cost of phototoxicity. We report that digital holo-tomographic microscopy (DHTM) reveals distinct patterns of virus infections in live cells with minimal perturbation. DHTM is label-free, and records the phase shift of low energy light passing through the specimen on a transparent surface. DHTM infers a 3-dimensional (3D) tomogram based on the refractive index (RI). By measuring RI and computing the refractive index gradient (RIG) values DHTM unveils on optical heterogeneity in cells upon virus infection. We find that vaccinia virus (VACV), herpes simplex virus (HSV) and rhinovirus (RV) infections progressively and distinctly increased RIG. VACV, but not HSV and RV infection induced oscillations of cell volume, while all three viruses altered cytoplasmic membrane dynamics, and induced apoptotic features akin to the chemical compound staurosporin, but with virus-specific signatures. In sum, we introduce DHTM for quantitative label-free microscopy in infection research, and uncover virus-type specific changes and CPE in living cells at minimal interference.

## Introduction

Viruses have a dual nature, the particle and the infected cell. At the onset of an infection the particle introduces proteins, DNA or RNA, and sometimes lipids into the host cell. The infected cell either produces viral components that are encoded in the viral genome, or raises an immune reaction against the virus, and silences the infection. In the former case, the infected cell develops a cytopathic effect (CPE). CPEs are diagnostic hallmarks of a particular virus, and well known to occur in cell cultures (for reviews, see (1, 2)). CPE can predict clinical outcomes *in vivo*. Examples include exacerbation of steatosis by hepatitis C virus, apoptosis in trigeminal ganglia by herpes simplex virus (HSV), or aseptic meningitis, paralysis, cardiomyelitis, and herpangina by enteroviruses, including poliovirus, coxsackievirus and enterovirus (EV) type 71 (3–5). While virus-induced CPE and cell death are exacerbated by cytokine responses, cytotoxic T cells or natural killer (NK) cells, virus-induced CPE of cultured cells apparently proceeds in a cell-autonomous manner.

The nature and the extent of CPE depend on the virus, the cell type, the host innate response, and the progression of infection. For example, distinct levels of CPE correlate with the amounts of newly synthesized virus particles (6). In case of adenovirus infected cells may lyse and release large amounts of viral particles following strong CPE cells, while persistently infected cells produce low amounts of progeny over long terms and have weak CPEs (7, 8). Yet, the extent of CPE not always correlates with virion production, as cells undergoing programmed cell death feature strong CPE at low viral titers (9–11). As viruses hijack cellular resources, CPEs elicited by virus infection may have distinct features, such as loss of membrane integrity, cell shrinkage, increased chromatin density, cell detachment from the substratum, formation of syncytia, loss or enforcement of the cytoskeleton, and reorganization of intracellular membranes (1, 12, 13). Despite the predictive nature of CPE for clinical and biological infections, time-resolved 3D analyses of virus-induced CPE are missing.

Here we describe a new approach using 3D digital holotomographic microscopy (DHTM) to study of the CPEs induced by three different viruses: vaccina virus (VACV), a large DNA virus replicating in the cytoplasm, herpes simplex virus type 1 (HSV-1), a large DNA virus replicating in the nucleus, and rhinovirus (RV), a small RNA virus replicating on cytoplasmic membranes. Classical video-enhanced contrast optical microscopy, such as interference microscopy and bright-field microscopy are limited by uneven image field intensity, lack of tomographic information and a focus-dependent size inflation of structures due to diffraction limitation (14). In contrast, DHTM allows prolonged quantitative time-resolved 3D image acquisition using an ultra-low powered laser (520 nm class 1 with 0.2 mW/mm^2^) without recognizable phototoxicity at high temporal and spatial resolution (15).

## Results

We infected HeLa cells with the VACV strain Western Reserve harboring an early/late GFP expressed transgene (VACV_WR E/L-GFP, in the following referred to as VACV-GFP) at a multiplicity of infection (MOI) of 2, fixed the cells 8 h post infection (pi) with para-formaldehyde, and recorded the refractive index (RI), nuclear DAPI stain and GFP intensity by correlative DHTM and fluorescence microscopy and compared to non-infected cells. At 8 h pi, all cells inoculated with VACV-GFP were infected based on their GFP intensity and DAPI staining of cell nuclei, whereas the uninfected cells’ GFP intensity was in the range of the background (Fig. 1A). Based on the RI change across a volume, the RI gradient (RIG) can be computed across the whole cell, similar to the RIG across an index gradient lens (for a simplistic illustration, see Fig. 1B, C).

**Fig. 1.**
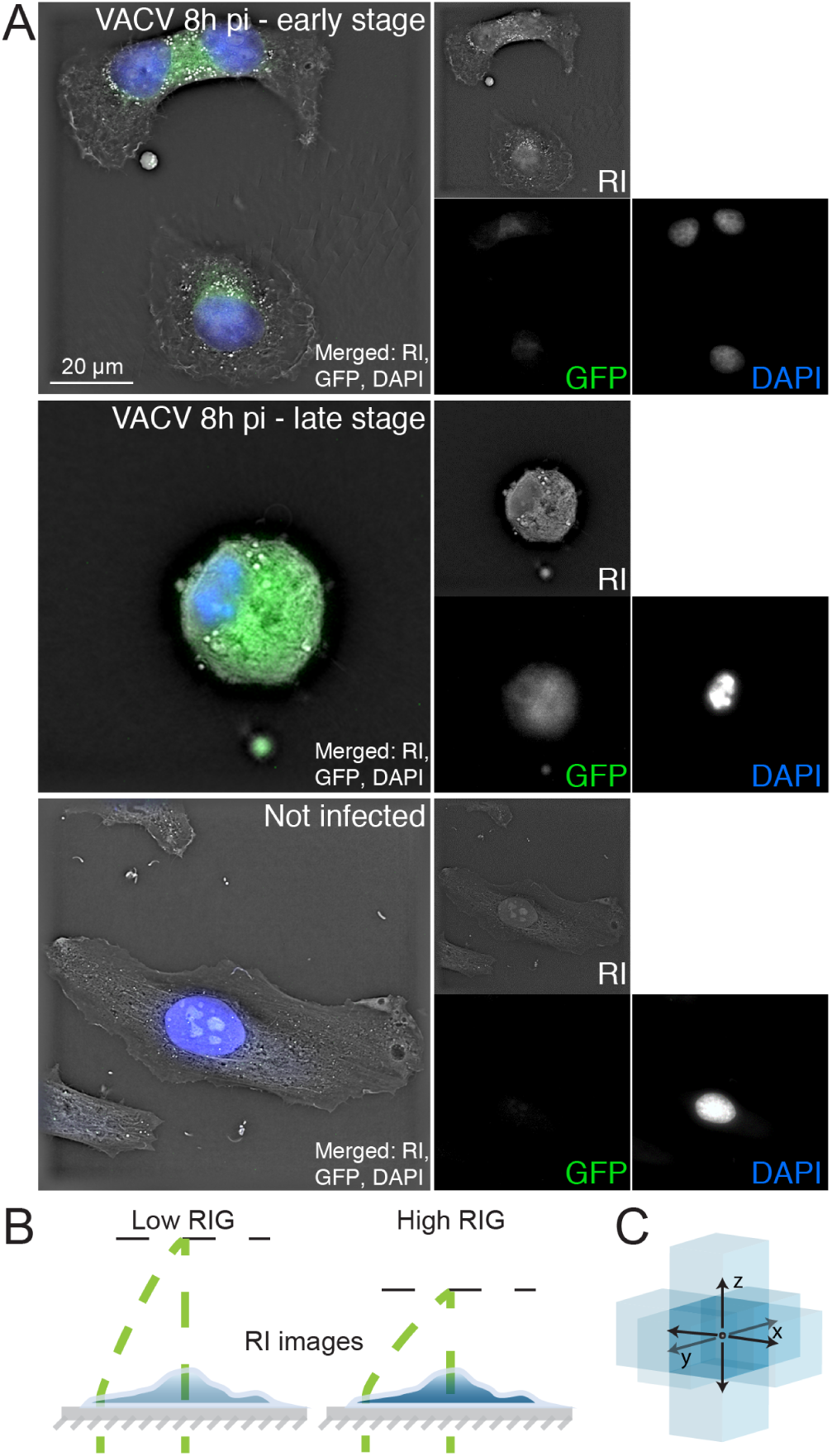
Correlative digital holo-tomographic microscopy and light microscopy of fixed and permeabilized cells. (A) Representative images of HeLa-ATCC cells incubated with 2 ml RPMI at 37° C for 8 h, fixed in 4% PFA in PBS, and imaged with a DHTM microscope and an attached epifluorescence module. Cells were either infected with VACV expressing GFP under control of early/late promoters (VACV_E/L-GFP; top and middle panel) at MOI 2, or remained uninfected (bottom panel). Top panel depicts a VACV-GFP-infected cell in early infection stage. Middle panel depicts rounding cell, indicating late stage VACV infection. Bottom panel depicts an uninfected cell. RI is displayed in grey, GFP in green, nuclei stained with DAPI appear in blue. Scale bar indicates 20 *µ* m. (B) Schematic illustrations of RI computation. A cell can be thought of as a gradient index microlens, changing its optical properties depending on biochemical activities. (C) RIG is derived from RI and represents a voxel-based measurement of the difference of the refractive index in 3D space. The RIG value of the voxel in the middle is represented as a middle blue box, which is calculated based on the difference to the light-blue blue voxels in the 3D neighborhood. Note that the reference beam (curved green dashed line) does not pass through the sample. RI is based on changes between the beam (straight green dashed line) and the reference beam.

We next imaged VACV-GFP infected and uninfected cells in live cell mode by DHTM at early to late stages of VACV infection in 2 h intervals up to 8 h pi (Fig. 2A). A progressive and prominent change in RI was observed in the infected cells, visualized in scaled pseudo-color. In contrast, the uninfected cells as well as the VACV-GFP infected cells treated with the deoxy-nucleoside analogue cytarabine (AraC), an inhibitor of VACV late gene expression (16, 17), showed less prominent RI changes, although AraC-treated cells inoculated with VACV-GFP exhibited a strong increase in RI in the cell nucleus. We quantified the RI change by deriving the RIG values across the entire cells (Fig. 2B). The RIG values of the VACV-GFP-infected cells gradually increased over the progression of the infection, reaching 3-fold at 8 h pi compared to the RIG before infection. In contrast, the RIG values of uninfected cells remained largely constant over 8 h. The AraC-treated cells showed only a small RIG increase of about 1.3-fold, suggesting that RIG increase is predominantly due to viral late gene expression. VACV infections exhibit membrane blebbing phenotypes at early and late time points, as well as focal bud-like swellings (18). Such features are diagnostic of a contractile cell cortex, as described in uninfected cells in cell migration and response to mechanical cues (19). We noticed an increased fraction of blebbing cells over 2-8 h pi in presence of AraC (Fig. 2C). This is distinct from membrane blebbing during entry (20), and suggests that blebbing at late infection stages does not require viral late gene expression, and does not affect the RIG.

**Fig. 2.**
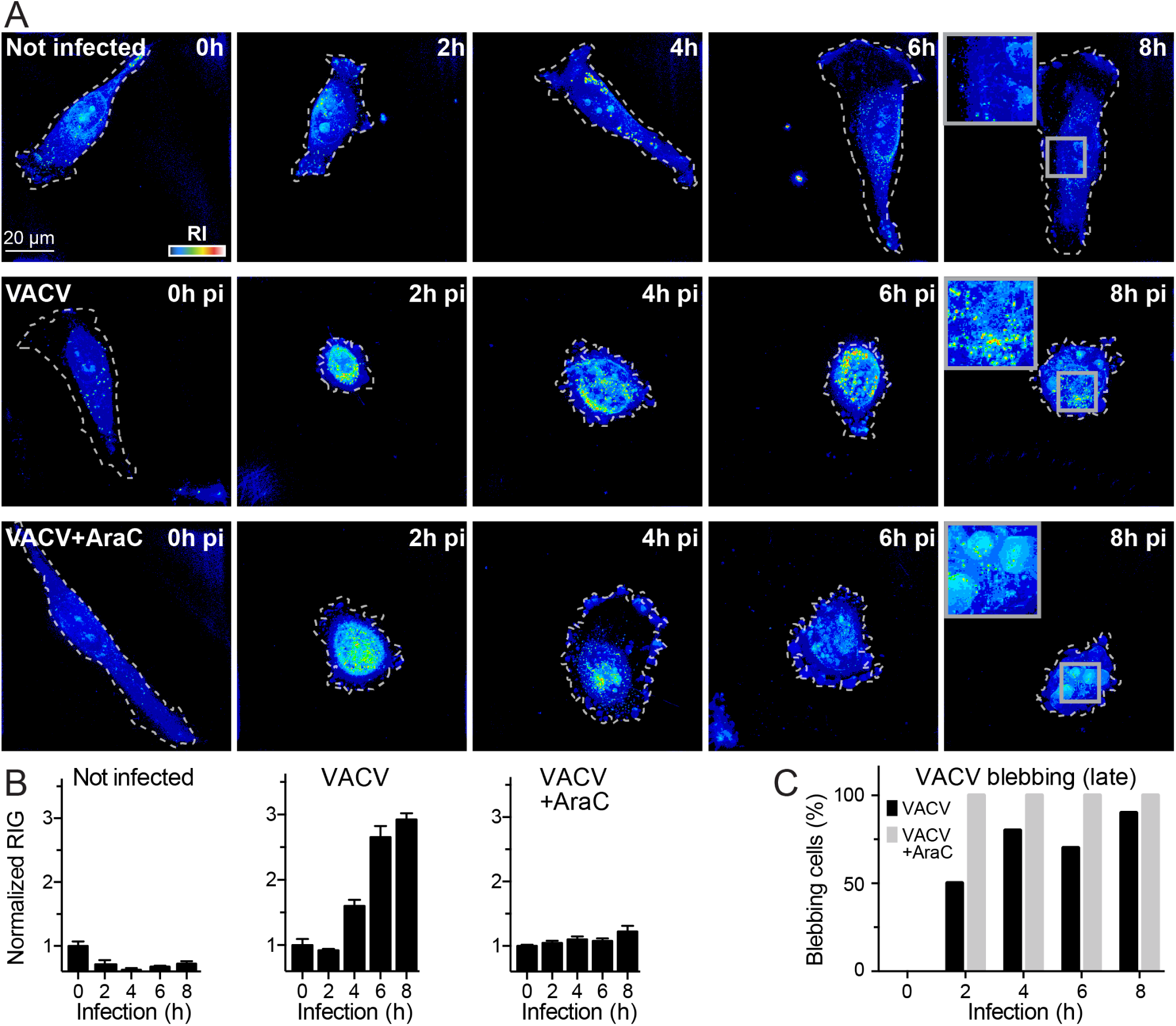
VACV late gene expression increases the cellular RIG. (A) Representative images of HeLa-ATCC cells incubated with 2 ml RPMI at 37° C for 8 h, and imaged at 2 h intervals by DHTM. Cells were uninfected, infected with VACV-GFP at MOI 2, or infected with VACV-GFP at MOI 2 in presence of 10 *µ* M AraC. Cell perimeters on the coverslip are outlined with white dashed lines. RIs are depicted as intensities values in a “thermal” lookup table. Images were obtained as holograms and depicted as projections of maxima along the z-axis of the 3D stacks. Scale bar indicates 20 *µ* m. (B) Cell RIG quantification of (A). RIG values were normalized to 1 for 0 h pi. Bars depict mean values *±* SEM of at least 10 cells for each condition and time point. (C) Comparison of the frequency of “blebbing” phenotype in VACV-GFP infected cells and VACV-GFP-infected cells in RPMI containing 10 *µ* M AraC. Data from at least 10 cells per time point and condition were acquired and manually scored as blebbing or non-blebbing.

To test if similar morphology changes of the VACV-GFP infected cells can be observed with an alternative imaging modality, we infected HeLa cells with VACV-GFP at MOI 2, and imaged cells in live phase-contrast and fluorescence modes at 5 min intervals for 8 h (Fig. 3, Video1.mp4) in a high-throughput light microscope. In agreement with the DHTM imaging results, the VACV-GFP infected cells expressed the GFP transgene at increasing intensity over the course of infection. As observed in the DHTM experiments, we observed host cell rounding, contractions and late stage blebbing of the infected cells, while the features of the uninfected cells remained largely invariant.

**Fig. 3.**
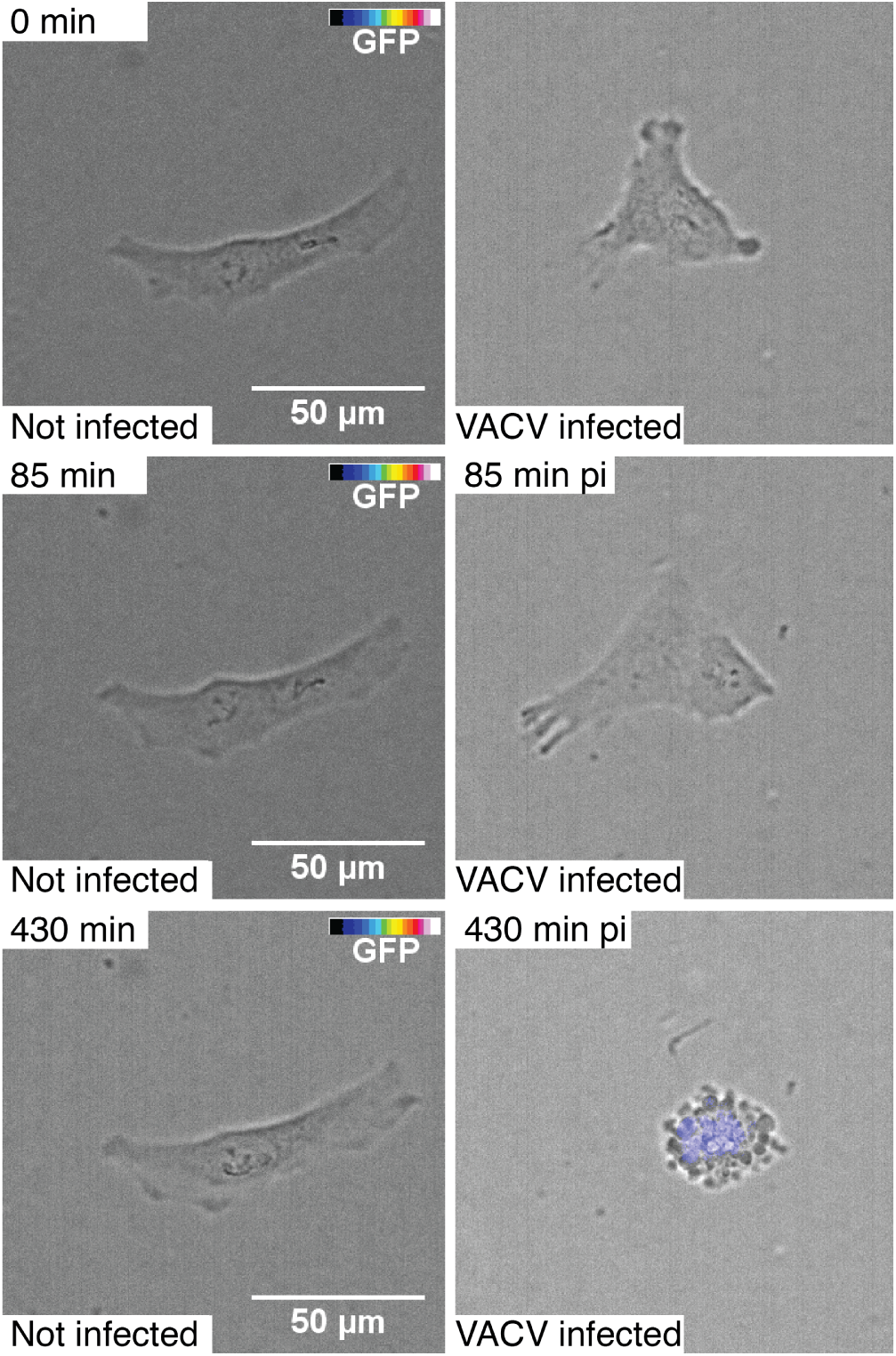
Cell morphology and VACV-GFP transgene expression visualized by automated and correlative phase-contrast and fluorescence live time-lapse microscopy. HeLa-ATCC cells were uninfected (left panel) or infected (right panel) with VACV-GFP using the cold binding protocol (30 min inoculation on ice, wash and transfer to 37° C). Cells were imaged with a high-throughput wide-field microscope every 5 min for 8 h. Cells were visualized with transmission light, the GFP fluorescence was intensity color-coded (color bar ranging from transparent through blue to white). Scale bars indicate 50 *µ* m. See also Video1.mp4

We next investigated the dynamic properties of the VACV-GFP-infected cells by using the holographic information, including cell volume. We reasoned that a change in cell volume might result in changes of the density of the cytoplasm, and hence affect the refractive properties of the cell. The accuracy of volumetric measurements based on DHTM was first assessed with polystyrene beads of different nominal sizes ranging from 0.5 to 4 *µ* m (Sup. Fig. 1A). Bead size was determined by two different analysis methods, a pixel-based method using the vendor’s software, STEVE, and an object-based 3D-surface segmentation method using Imaris (Sup. Fig. 1B). Both procedures yielded similar results, although the object-based method were slightly more accurate.

To measure cell volume by DHTM, we used the image stack segmentation procedure in Imaris. Time-resolved 3D live cell DHTM imaging of infected and uninfected cells between 0 and 8 h pi indicated that the VACV-GFP infected cells underwent a series of shrinkage and dilation phases, while the volume of the uninfected cells remained largely constant (Fig. 4, Video2.mp4, Video3.mp4). The first shrinkage phase peaked at about 100 min pi and reduced the cell volume by about 50%. This contraction was followed by a steady recovery phase restoring the original volume at about 180 min pi. The onset of the next shrinkage period was at 230 min, and lasted about 40 min, followed by a short recovery phase of about 30 min, and a period of stable volume with minor fluctuations until 440 min pi, when another shrinkage period occurred. We did not detect a correlation between cell shrinkage or expansion with the RIG values, the latter steadily increasing over the course of infection. We conclude that the VACV-GFP infection-induced RIG increase is not caused by generic changes of the cell volume. To further explore the correlations of RIG and virus-induced CPE, we recorded RIG in cells infected with HSV-1-GFP harboring a GFP transgene under the constitutively active CMV promoter, and RV-A1a at 0, 8, 10, 12, 14 and 16 h pi (Fig. 5A). In HSV-1-GFP-infected cells, CPE was observed from 8 to 10 h pi on, followed by cell surface roughening at 16 h pi. Remarkably, RIG did not increase despite of the onset of CPE prior to 16 h pi (Fig. 5B). Cells infected with RV-A1a showed strong CPE starting from 10 and 12 h pi, involving the condensation of cytoplasm. The RIG of RV-A1a infected cells steadily increased at 10 to 16 h pi up to about 2.5-fold of the RIG of the uninfected controls. This correlated with the induction of apoptosis, virus-controlled necroptosis, and the loss of cytoskeletal elements, such as F-actin, as observed in picornavirus infections (21–23). Remarkably, RV-A1a-infected cells adopted a transient branched shape at 12 to 14 h pi, before rounding up at 16 h pi (Fig. 5A).

**Fig. 4.**
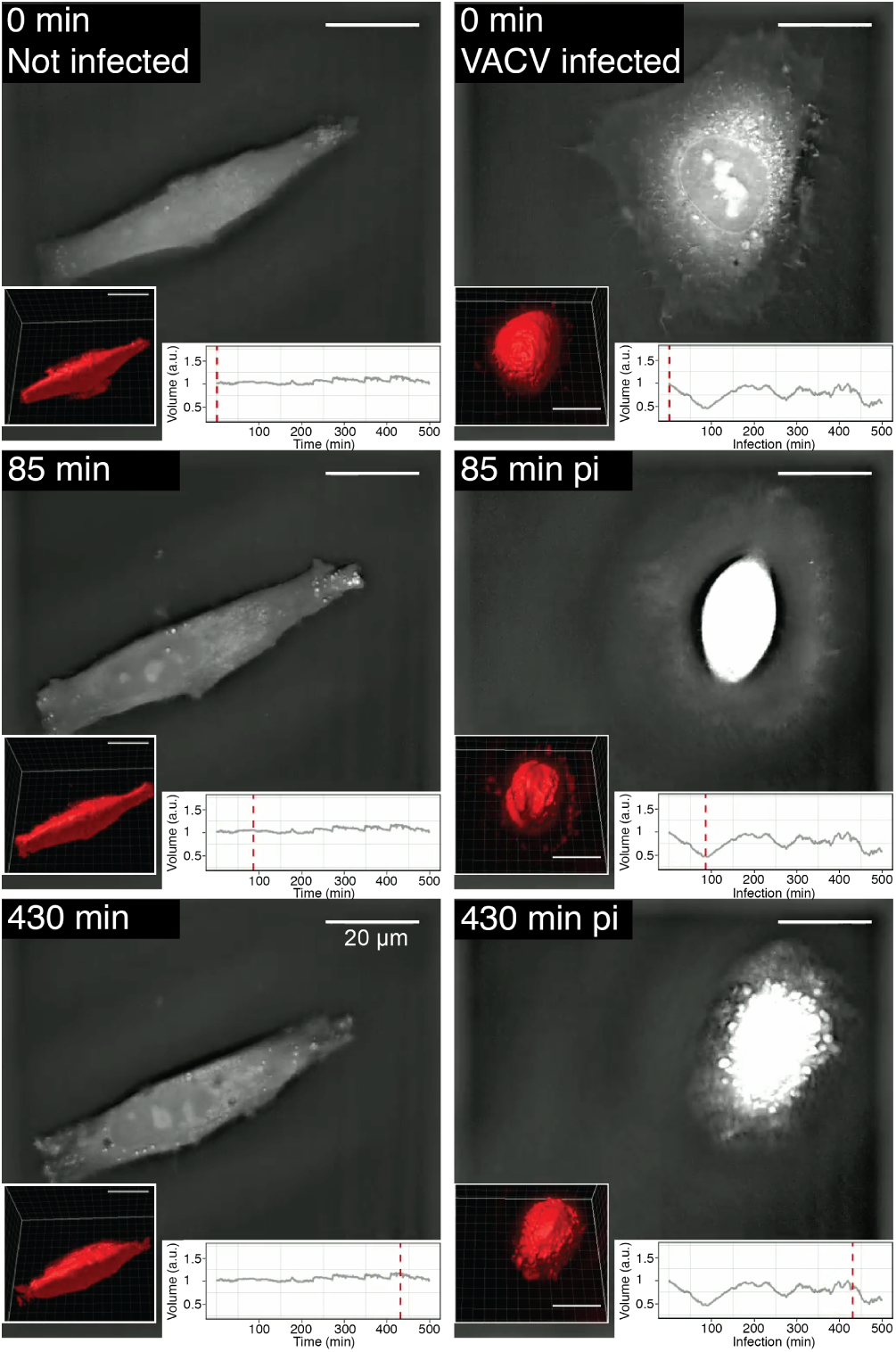
Cell morphology and volume dynamics of VACV-GFP-infected cells, visualized by label-free time-lapse DHTM. HeLa-ATCC cells were infected by VACV-GFP following the cold binding protocol. Cell holograms were acquired every minute for 8 h, RI shown as greyscale images. Volume measurement was performed using Imaris software by surfaced fitting and 3D rendering (see the lower left corner of each frame). Plots show the relative volume normalized to 0 min pi. The red dashed lines correspond to the time points related to the corresponding hologram. Scale bars indicate 20 *µ* m. See also Video2.mp4, Video3.mp4.

**Fig. 5.**
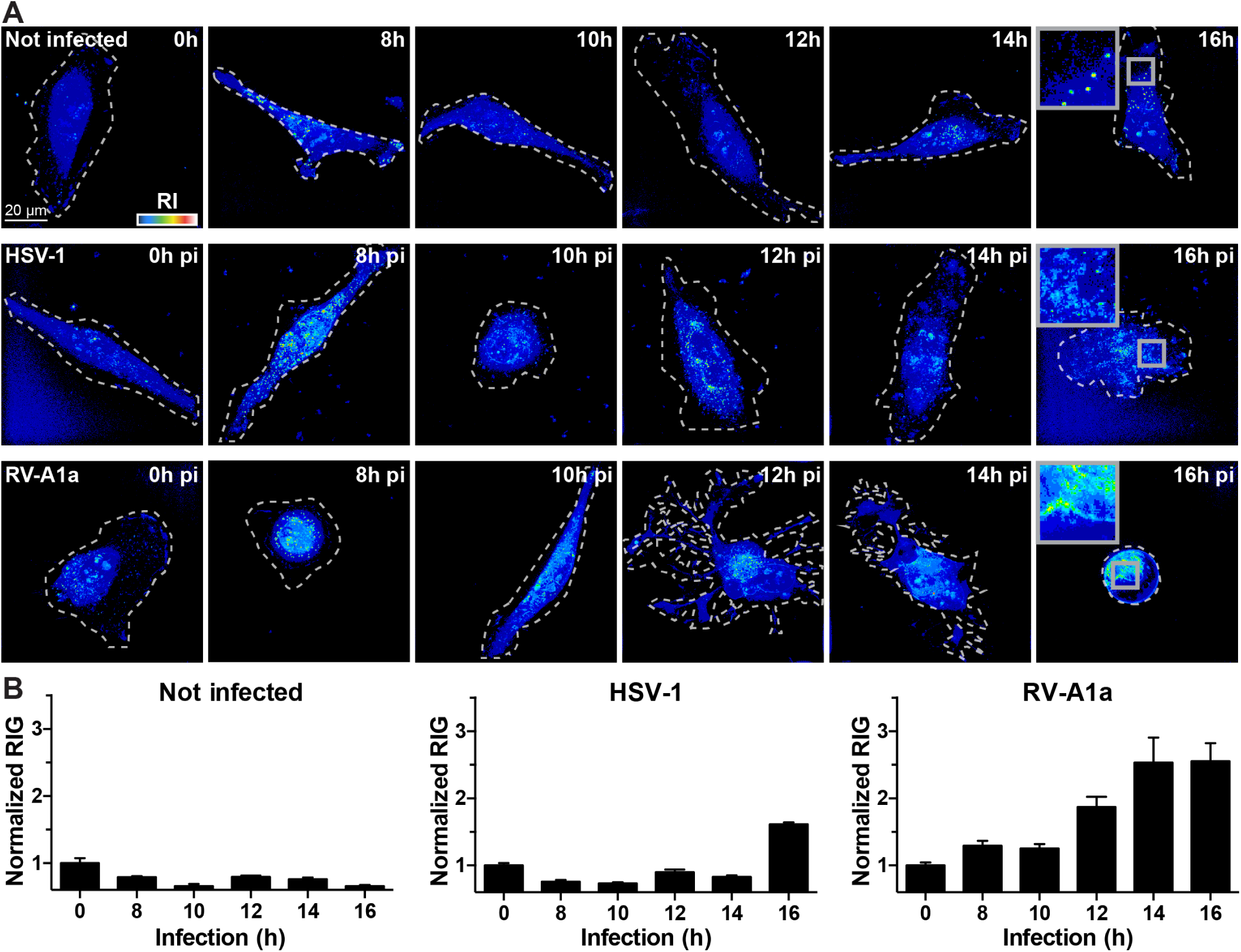
HSV-1 and RV increase cellular RIG late in infection. (A) Representative images of uninfected, HSV-1-GFP (MOI 10) infected HeLa-ATCC cells or RV-A1a (MOI 50) infected HeLa Ohio cells, imaged by a DHTM microscope at indicated time points. Cell perimeters on the coverslip are outlined with white dashed lines, and the refractive indices are depicted as intensities in a “thermal” lookup table. Images were obtained as holograms, and the 3D stacks depicted as z-projections of maxima. Scale bars indicate 20 *µ* m. (B) Normalized cell RIG of the data shown in (A). Bars depict mean values *±* SEM of at least 10 cells for each condition and time point.

Two distinct inhibitors of RV replication were employed to test if the RIG increase was directly associated with RV-A1a replication. The first inhibitor, PIK-93, blocks phosphatidylinositol-4-kinase class 3beta (PI4K3b) activity, and thereby precludes the lipid remodeling by counter-current lipid fluxes for virus replication (24). In absence of PI4K3b activity, viral replication is suppressed, because the altered lipid composition of the replication membrane can no longer support viral host proteins necessary for virus replication (reviewed in (25)). The second inhibitor, MLN9708, is a second generation inhibitor of the proteasome, which is required for enterovirus replication (26–28). Both PIK-93 and MLN9807 blocked the appearance of high RI/RIG structures, although the effect of PIK-93 was incomplete (Fig. 6). The partial effect of PIK-93 on RIG was, however, consistent with the partial reduction of PI4P levels and the incomplete inhibition of RV-A1a replication (24, 29). These results support the notion that an increase of RIG in virus-infected host cells occurs in a virus type-specific manner, and that RIG changes are a reliable indicator of early and late CPE. The latter is often associated with virus-induced apoptosis, necrosis, or necroptosis. We therefore tested if the apoptosis-inducing agent staurosporine (30) also induced a RIG response. DHTM tomography of 10 *µ* M staurosporine-treated cells (Fig. 7A) confirmed that staurosporine induces an apoptotic phenotype as early as 1 h post treatment. Image quantifications indicated that RIG increased up to 4 h post application, and slightly decreased at 5 h (Fig. 7B). The extent of the RIG increase reached similar levels as upon RV-A1a infection, thereby confirming that apoptotic phenotypes are tractable by DHTM.

**Fig. 6.**
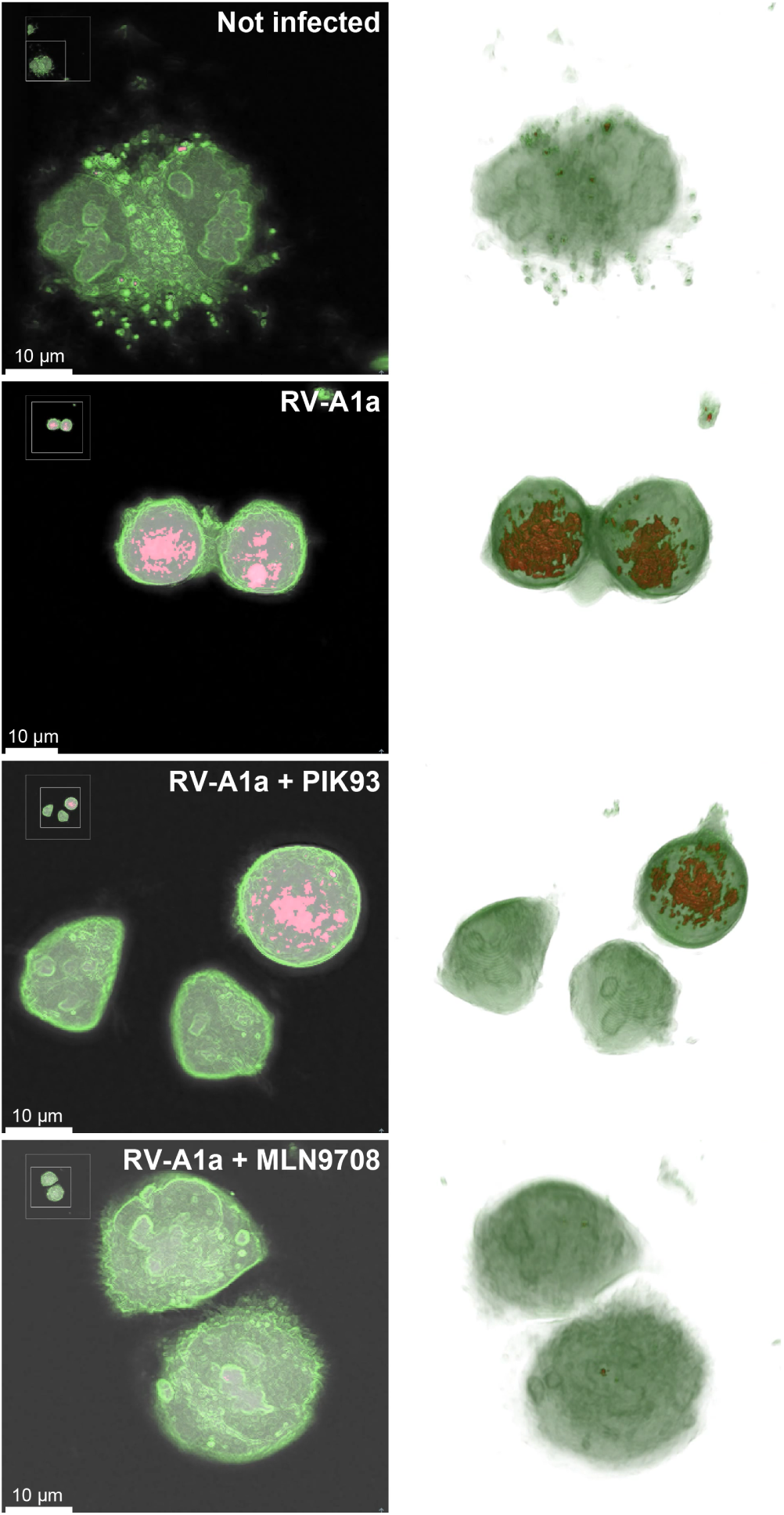
Drug treatment inhibits changes in RIs of RV-A1a-infected cells. Representative images of mock infected (top panel), or RV-A1a-infected (MOI 50) HeLa Ohio cells (lower three panels). Cells were treated with either PIK93 (5 *µ* M), or MLN9708 (10 *µ* M), and imaged by DHTM for 8 h at 1 min intervals. Cell membranes are labelled in green, high RI and RIG regions in red. The left row depicts central z-slices of reconstructed holograms, and the right row 3D reconstructions of the holograms. See also Videos4-7.mp4. Scale bars as indicated.

**Fig. 7.**
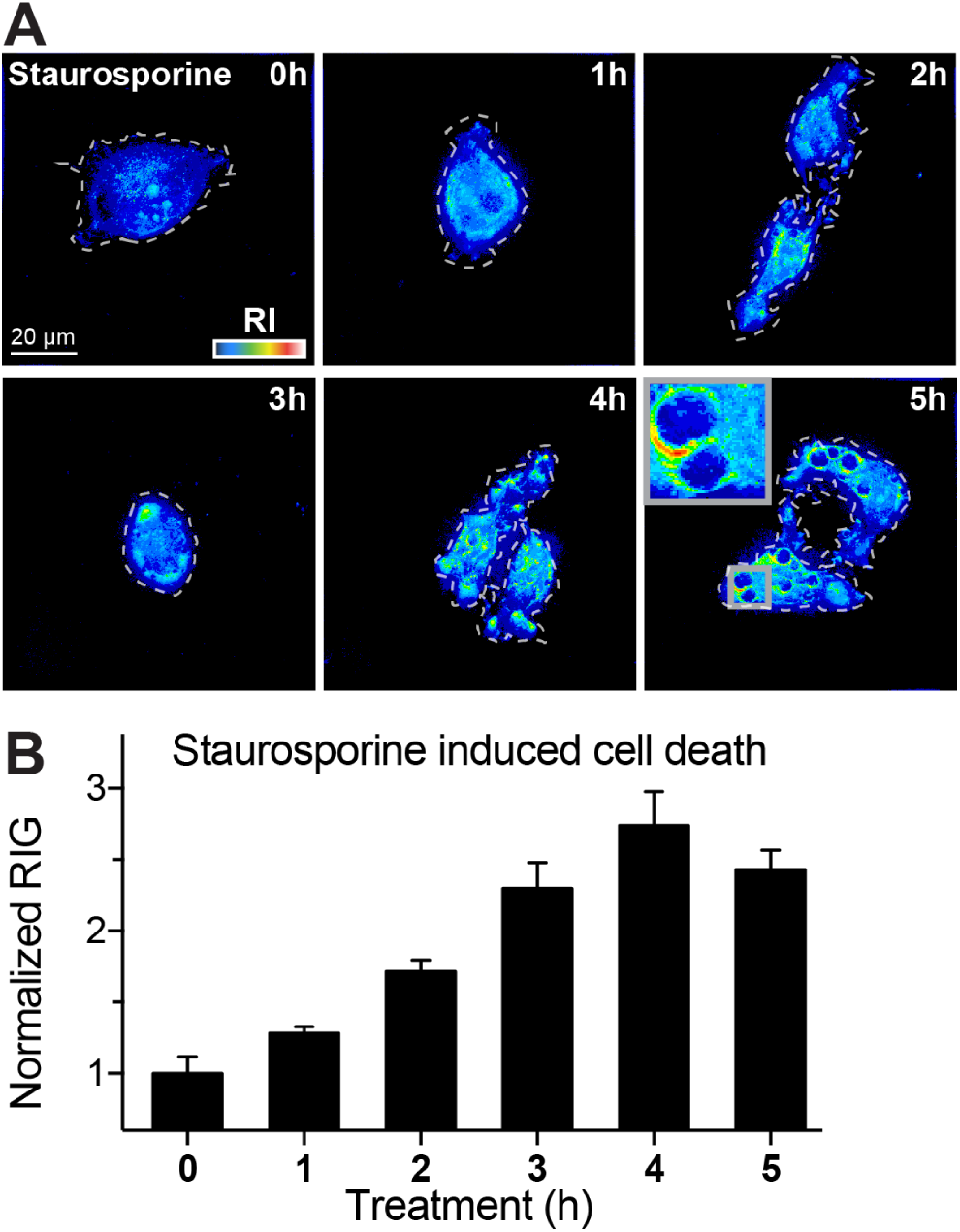
Staurosporine-induced apoptosis rapidly increases the cellular RIG. (A) Representative DHTM images of HeLa-Ohio cells incubated with staurosporine (10 *µ* M) at 37° C for 5 h. Cell perimeters on the glass coverslip are outlined with white dashed lines, and RI depicted as intensities in “thermal” lookup table. Images were obtained as holograms, and the 3D stacks depicted as z-projections of maxima. Scale bar indicate 20 *µ* m. (B) Cell RIG quantification of (A). RIG values were normalized to 1 at 0 h post addition. Bars depict mean values *±* SEM of at least 10 cells for each condition and time.

## Discussion

DHTM is a noninvasive label-free light microscopy technology. It measures the RI of a transparent object by an interference procedure (31–36). DHTM is superior to classical optical techniques, such as tomographic phase microscopy and diffraction phase microscopy (37), because of its high sensitivity, accuracy and non-invasiveness (15). The system produces quantitative 3D information at high spatial and temporal resolution. It readily measures the volume of cells and subcellular structures. Furthermore, it can be used to acquire two new dimensions of cell data, RI and RIG, which cannot be readily assessed by other imaging modalities. The methodology is robust, and overcomes a series of technical limitations in live cell pathogen imaging, including the labeling of cellular and viral entities by chemical or genetically-encoded fluorophores, and toxicity ensuing imaging (for recent reviews, see (38, 39)). The non-invasiveness and low laser power required for DHTM underscore the suitability of DHTM for long-term live imaging.

Here, we introduce DHTM to virus research by employing three different viruses, VACV of the *poxviridae*, HSV-1 of the *herpesviridae* and RV from the *picornaviruses*. VACV is an enveloped, double-stranded DNA virus, replicating and assembling particles in the cytoplasm (40). It is used to immunize and protect humans against small pox caused by variola virus, one of the deadliest viruses to man (41, 42). VACV exploits apoptotic mimicry to enter host cells through cellular blebbing, that is, protrusions of the cell membrane, implicated in cytokinesis and cell motility (20, 43, 44). VACV-induced morphological changes of host cells include cell rounding, blebbing and bud-like swellings (18). Picornaviruses are small, positive-sense single-stranded RNA viruses with a non-enveloped icosahedral capsid of 28-30 nm in diameter (45). RVs replicate in the cytoplasm on Golgi-derived membranes that are closely associated with the ER, and elicit a strong CPE in tissue culture (21, 24, 25). HSV-1 is an enveloped, double-stranded DNA virus, well adapted to human hosts and controlled by innate immunity including the interferon induced human myxovirus resistance protein B (MxB) (46). If HSV-1 breaks through the innate host defense, it replicates in the cell nucleus and sheds progeny prior to cell lysis (47, 48). In its latent state, HSV-1 evades the immune system, avoids CPE and downregulates apoptosis (49, 50). Upon reactivation, HSV-1 is transmitted to the epidermal tissue where it causes lytic infection manifested as epidermal blisters. We initially used DHTM to determine the volume of cells, and became aware of periodic shrinkage and expansion of VACV-infected cells, as well as membrane blebbing at late stages of VACV but not in RV-A1a infections. Cell volume dysregulation is known to contribute to disorders, such as liver insufficiency, diabetic ketoacidosis, hypercatabolism, fibrosing disease, and sickle cell anemia, its regulation affects cell proliferation, and apoptosis (51).

VACV infection-induced cell volume oscillation implies that VACV regulates membrane trafficking, such as endocytosis and secretion, and possibly also ion channels. For example, the release of potassium, chloride and bicarbonate ions is known to trigger cell shrinkage, or the accumulation of sodium, potassium and chloride ions cell swelling through the activation of cotransporters, exchangers or channel proteins (51). In addition, shrinkage and swelling processes involve organic osmolytes, such as sorbitol and glycerophosphorylcholine, which accumulate in shrinking cells and are released in swelling cells. How exactly VACV controls these processes to produce progeny remains an open question.

In addition to the accurate, noninvasive readout of cell volume, the RIGs measured by DHTM have served as an indicator for the granularity of cell structures (52). We show that specific RIG signatures indicate virus-induced CPEs. CPE follows cell shrinkage and blebbing, and terminates by cell death (13, 53). In fact, viruses control cell death processes, including apoptosis or necroptosis, for example by diverting upstream regulating kinases, such as RIPK1 in RV-infected cells (2, 21). Other viruses prevent infection-induced apoptosis at early stages, gaining crucial time for production of virus progeny (50, 54–56). At late infection stages, viruses gain virulence by enhancing progeny virus release from the cells (56–58). This leads to the notion that lytic infection resembles a necrosis-like program, leading to the release of both cellular contents and virus particles (7, 59).

CPE are elicited not only by wide range of pathogens, including viruses and bacteria, but also emerge in malignant cells during transformation, and in immune defense by cytotoxic cells for instance (6). Despite such insights, the CPE has remained one of the least understood processes in infection biology and pathology. This is in part a plethora of virus-induced cell reactions tune the infection, including pro- and anti-viral signaling and innate immunity response. CPE arises because of the many altered processes in an infected cell. It is triggered by a handful of viral genes introduced into a naïve cell, and comprises pro- and anti-viral components. Our results show that all tested viruses VACV, HSV-1 and RV-A1a induced RIG increase over the course of infection, with virus-specific kinetics, extent and signature. Importantly, the RIG measurements scored CPE independent of cell contractions and blebbing, which can not be uncoupled in phase contrast microscopy. This indicates that RIG truly measures new dimensions in CPE that have not been accessible with conventional microscopy analyses. It underlines the superiority of DHTM over phase contrast microscopy, and allows the attribution of virus-specific processes to CPE. In case of VACV infection the RIG increase appeared to require late viral gene expression, since RIG changes were impaired by AraC, which inhibits late but not intermediate or early genes. For RV, we infer that at the onset of RIG increase at 8 h pi not only cytoplasmic membranes are being rearranged and expanded for viral replication, but also phase separation processes occur in the perinuclear area, a hallmark of RV-infected cells. With HSV-1, RIG increase was most dramatic at 16 h pi, with large changes in the nucleus and the cytoplasm, indicative of processes associated with virion production and release from the infected cell.

In conclusion, we show that DHTM is suitable to analyze cell biological processes in virus-infected cells at high spatiotemporal resolution, without the need to introduce cell markers, or using high intensity laser light. DHTM convinces with high spatial resolution with potential super-resolution options (32, 60). It enhances insights into CPE, a complex process induced by nearly all pathogens, and increasingly mined by high-throughput screening assays (61). DHTM promotes the dynamic analyses of virus assembly, for example in phase separated zones of the cytoplasm and the nucleus, and provides a deeper understanding of the nature of viruses.

## ACKNOWLEDGEMENTS

We thank all members of the Greber lab for discussions and comments throughout the course of this work. We thank Nicole Meili and Melanie Grove for tissue culture and general lab support. We acknowledge excellent technical support from Dr. Lisa Pollaro, Luca Clario and Dr. Sorin Pop (Nanolive SA, Ecublens, Switzerland). We thank the Center for Microscopy and Image Analysis at the University of Zurich headed by Dr. Urs Ziegler for providing polystyrene beads. We thank Dr. Jason Mercer (MRC LMCB, University College London, London, United Kingdom) for kindly providing Vaccinia virus IHD-J E/L-GFP. UFG acknowledges financial support from the University of Zurich, the Swiss initiative in Systems Biology SystemsX.ch (VirX), and the Swiss National Science Foundation (grant numbers 310030B_160316 and 31003A_179256 / 1).

## Materials and Methods

### Cell lines and viruses

HeLa-ATCC from the American Type Culture Collection (ATCC) were maintained in Dulbecco Modified Eagle Medium (DMEM; GIBCO-BRL) containing 10% fetal calf serum (FCS), non-essential amino acids (NEAA), and penicillin-streptomycin (GIBCO-BRL) at 37° C and 5% CO_2_. All cell cultures were maintained in a cell bank system and kept in low passages for all experiments. Vaccinia virus strain International Health Department J (VACV-IHD-J) containing early / late (E/L) GFP transgene was kindly provided by J. Mercer (University College London, UK) (17, 62). To obtain the purified mature virions (MVs) cytoplasmic lysates were pelleted through a 36% sucrose cushion for 90 min with Optima XPN-100 ultracentrifuge (Beckmann Coulter) SW32Ti rotor at 18,000 rpm. The viral pellet was suspended in 10 mM Tris pH 9.0, and virus separated from contaminating material on a 25 to 40% sucrose gradient at 14,000 rpm for 45 min. Following centrifugation, the viral band was collected by aspiration and concentrated by pelleting at 14,000 rpm for 45 min. MVs were suspended in 1 mM Tris-HCl pH 9.0 and titered for plaque forming units (PFU) per ml as previously described (63). The above method revealed an infectious particle to PFU ratio of VACV-WR and VACV-IHD-J virus on monkey kidney BSC40 cells in the range of 50:1 and 80:1, respectively. RVs were grown in HeLa cells as described (64, 65). Cells were inoculated with a lysate from infected cells at 33.5° C overnight. When CPE was visible in 80-90% of the cells, the media was removed and cells harvested by scraping and pelleting. Cells were lysed by three freeze/thaw cycles in liquid nitrogen followed by addition of 1% NP40 and homogenization with a Dounce homogenizer. The suspension was centrifuged at 2,500 *×* g for 10 min and the supernatant transferred into a new tube. Free RNA was digested by addition of 150 *µ* g RNase per 10 ml and incubation at 37° C for 30 min. Virus was purified on a CsCl gradient and extensively dialyzed against 140 mM NaCl, 25 mM Hepes, 5 mM MgCl_2_. Aliquots were stored at −80° C. The HSV-1 recombinant strain C12 expressing GFP from the major CMV promoter was kindly provided by S. Efstathiou (University of Cambridge, Cambridge, UK), and used as described (46, 66).

### Cold-synchronized infections

Media and PBS were cooled to 4° C to prevent endocytosis. Cold virus inoculum in HEPES-buffered RPMI (Sigma), containing 10% FCS, NEAA and penicillin-streptomycin (GIBCO-BRL) was added to HeLa cells and incubated on ice for 30 min. Next, cells were washed three times with cold phosphate buffered saline (PBS) and overlaid with either 37° C warm carbonate buffered DMEM (GIBCO-BRL) containing 10% FCS, NEAA and penicillin-streptomycin or 37° C warm HEPES buffered RPMI (Sigma) containing 10% FCS, non-essential amino acids and penicillin-streptomycin for automated microscopy or tomographic holography respectively.

### Automated time-lapse, multi-site, multi-channel microscopy

Time-lapse multisite, multichannel microscopy on HeLa cells grown in 96-well imaging plates (Greiner Bio-One) was performed with an ImageXpress Micro Widefield High-Content Analysis System (IXM-XL, Molecular Devices) microscope with a synthetic air to CO_2_ mixture of 95% to 5%, respectively, in a humidified environment at 37° C with a 20x objective.

### Live label-free holographic tomography

Live holographic tomography was performed using a 3D Cell Explorer microscope (Nanolive SA, Ecublens, Switzerland). Cells were grown and imaged using 35 mm Ibidi glass bottom *µ* -Dish dishes (Ibidi GmbH, Germany). During imaging, temperature (37° C) and humidity was controlled using an Ibidi Heating & Incubation System (Ibidi GmbH, Germany), while the pH was maintained by using RPMI containing 20 mM HEPES buffer.

### Cell tracking and volume measurement

To measure the cell volume, 3D stacks obtained by DHTM were digitally stained and voxel-segmented using STEVE (Nanolive SA, Ecublens, Switzerland), exported as TIF-files and imported into Imaris (Bitplane AG, Switzerland). Next, a surface was fitted to the imported 3D voxels aimed at complete volume segmentation. The fitted surface was tracked and enclosed features (volume, centroid position, centroid speed) were measured with Imaris over the entire duration of the time-lapse experiment. Finally, the volume of each cell in a time-lapse series was normalized to the cell volume at time zero.

To benchmark the volume measurements from holographic tomography, we used polystyrene beads of 0.5, 0.75, 1 (Fluoresbrite, Polysciences) and 4 *µ* m diameter (Tetraspeck, ThermoFisher). The beads were diluted in PBS, allowed to sediment to the bottom of the dish and imaged. We next quantified their volume either by voxel-segmentation and counting in STEVE or by surface fitting in Imaris (Sup. Fig. 1).

### Cell refractive index gradient measurement

RI of a material is defined as speed of light in vacuum divided by the speed of light in the particular material. RIG is a computed value describing RI change within a neighborhood of a particular voxel according to the Eq. 1.

Measurement of the relative RIG of the cells was performed by ‘digital staining’ in STEVE according to the user manual. The staining was aimed at segmenting at least 95% of the manually determined cellular signal; next, the mean RIG of the digital staining was calculated. Boundaries were adjusted by shifting the RI outside of the 95% constraint. Finally RIG of each cell in a time-lapse was normalized to the RIG of the cell at the start of the time-lapse.

## Equations

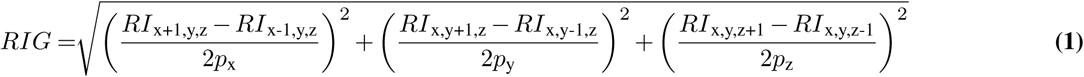

here 2*p*_x_, 2*p*_y_, 2*p*_z_ is the resolution of the image, and *RI*_x ± 1, y ± 1, z ± 1_ is the neighbouring pixels RI value.

## Appendix 1. Supplementary Information

This is a supplementary section to the preprint manuscript by Yakimovich & Witte et al. Supplementary Videos can be downloaded alongside with the manuscript. Source code used in this manuscript can be obtained from https://github.com/ayakimovich/DHMViruses. All further information is available upon request.

### Supplementary Figures

**Fig. S1:**
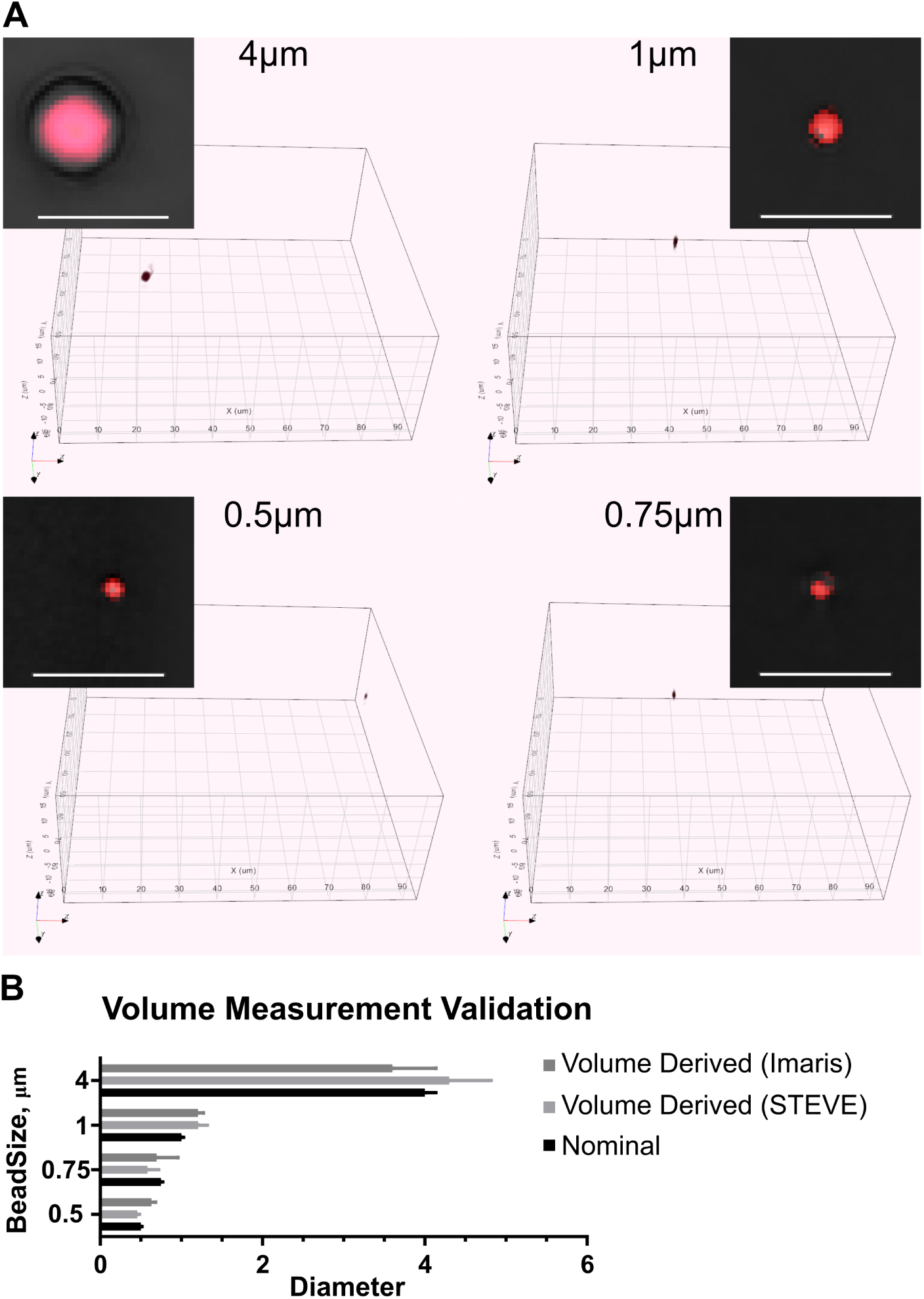
the volume measurement using polystyrene beads of defined size. (A) Polystyrene beads (Tetraspeck, ThermoFisher) of defined diameter were diluted in PBS, allowed to sediment to the bottom of the dishes and imaged using a 3D Cell Explorer microscope. Small boxes depict artificial RI stain of beads in red on black background. Large box shows full 3D visualization of acquired holograph. 3D stacks of the beads were digitally stained (voxel segmented) for an RI estimated to cover at least 95% of the bead volumes. Scale bars indicate 5 *µ* m. (B) Comparison of volume quantification performed by voxel summation either by voxel-segmentation and counting in STEVE software or by surface fitting in Imaris. Bead diameter was computed based on voxel counts and voxel size and compared to the nominal diameter provided by the manufacturer.

### Supplementary Videos

**Video 1: Cell morphology and VACV E/L-GFP transgene expression visualized by automated and correlative phasecontrast and fluorescence live time-lapse microscopy.**

HeLa-ATCC cells were either mock infected (left panel) or infected (right panel) with VACV_E/L-GFP virus (MOI 2). Cells were imaged with a high-throughput widefield microscope every 5 min for 8 h. Samples were visualized with transmission light. The GFP fluorescence was color-coded (color bar corresponding to fluorescence intensity from transparent through blue to white). Scale bar indicates 50 *µ* m. The video is related to Fig. 3.

**Video 2: Cell morphology and volume dynamics of an uninfected cell visualized using label-free time-lapse holographic tomography.**

HeLa-ATCC cells were treated with cold binding medium, followed by transfer to 37° C. Cell holograms were acquired every minute for 8 h and shown as greyscale images. Volume measurement was performed using Imaris software by surfaced fitting (3D rendering in the lower left corner of each frame). The volumes relative to 0 min time point are plotted. The red dashed lines depict the corresponding time points in volume plot. Scale bar indicates 20 *µ* m. The video is related to Fig. 4.

**Video 3: Cell morphology and volume dynamics of VACV infected cell visualized using label-free time-lapse holographic tomography.**

HeLa-ATCC cells were infected with VACV_E/L-GFP virus (MOI 2) by cold synchronization. Cell holograms were acquired every minute for 8 h and shown as greyscale images. Volume measurement was performed using Imaris software by surfaced fitting (3D rendering in the lower left corner of each frame), volume relative to 0 min is plotted. The red dashed lines depict the corresponding time points in volume plot. Scale bar indicates 20 *µ* m. The video is related to Fig. 4.

**Videos 4-7: Drug treatment inhibits RV-A1a induced RI changes**

HeLa-Ohio cells were left uninfected (top panel), or infected with RV-A1a (MOI 50, lower three panels). Cells were left untreated (upper two panels), or were treated with either 5 *µ* M PIK93, or 10 *µ* M MLN9708. Holographic images were acquired at 1 min intervals for 8 h. Cell membranes are labelled in green, high refractive index and refractive index gradient regions are labelled in red. Scale bars indicate 10 *µ* m. Still frames of the videos are provided in Fig. 6.

